# Reanalysis shows the extreme decline effect does not exist in fish ocean acidification studies

**DOI:** 10.1101/2022.04.11.487807

**Authors:** Philip L Munday

## Abstract

A meta-analysis published in PLoS Biology by Clements et al. (2022) claims there is an extreme decline effect in studies published between 2009-2019 on the impacts of ocean acidification (OA) on fish behaviour. Here I show that the extreme decline effect reported by Clements et al. is a statistical artifact caused by the way they corrected for zero values in percentage data, which was more common in the earliest experiments compared with later studies. Furthermore, selective choices for excluding or including data, along with serious errors in the compilation of data and missing studies with strong effects, weakened the effect sizes reported for papers after 2010, further exacerbating the decline effect reported by Clements et al. When the data is reanalyzed using appropriate corrections for zero values in percentage and proportional data, and using a complete, corrected and properly screened data set, the extreme decline effect reported by Clements et al. no longer exists.

## MAIN TEXT

A meta-analysis published in PLoS Biology by Clements et al. [1] claims there is an extreme decline effect in studies published between 2009-2019 on the impacts of ocean acidification (OA) on fish behaviour, with the modelled average effect size declining an order of magnitude, from >5 in 2009-2010 to <0.5 after 2015. Here I show that the extreme decline effect reported by Clements et al. is a statistical artifact caused by the way they corrected for zero values in percentage data, which was more common in the earliest experiments compared with later studies. Furthermore, selective choices for excluding or including data, along with serious errors in the compilation of data and missing studies with strong effects, weakened the effect sizes reported for papers after 2010, further exacerbating the decline effect reported by Clements et al. When the data is reanalyzed using appropriate corrections for zero values in percentage and proportional data, and using a complete, corrected and properly screened data set, the extreme decline effect reported by Clements et al. (Fig 1a,b) no longer exists. Instead, there is a more gentle and consistent decline in effect size magnitude through time (Fig 1c), from an average around 2 in 2009-2010 and remaining well above zero in 2018-2019 (Fig 1d).

**Figure 1:**
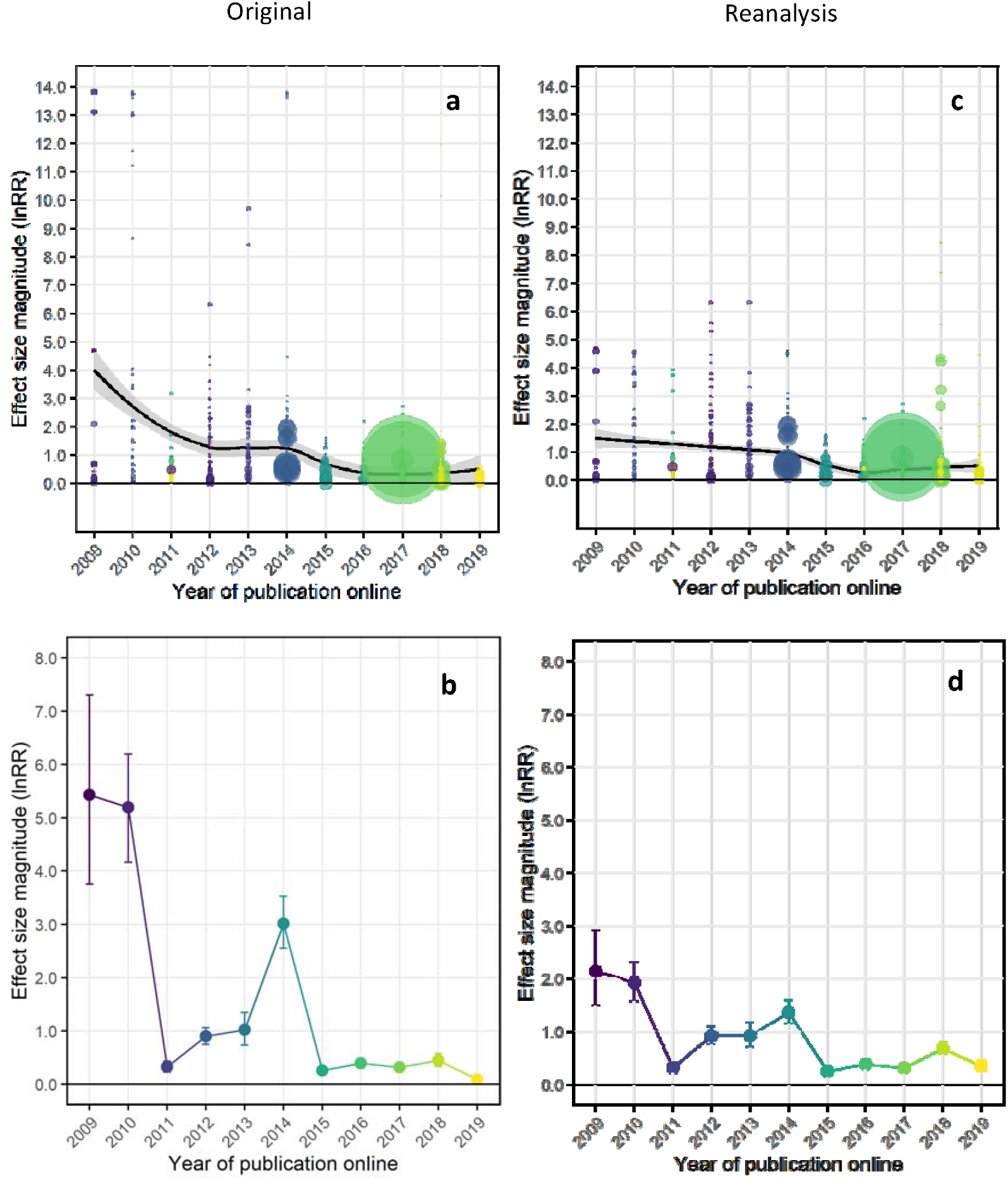
Original analysis of effect sizes in studies on the impacts of ocean acidification on fish behaviour by Clements et al. using 0.0001 to replace zero values in percentage and proportional data (a,b) and reanalysis with the corrected, updated and screened data set (supplementary data 1) using 1 to replace zero values in percentage data and 0.01 to replace zero values in proportional data (c,d). Top row shows all calculated effect sizes (lnRR) fitted with a Loess curve and 95% confidence bounds. Bottom row are modelled variance-weighted average effect sizes by year. Experiments with smaller variance are given greater weight in calculating the model means in the bottom row.

The primary reason for the extreme decline effect reported by Clements et al. is their decision to replace zero values in percentage data (range 0-100%) with a tiny value to four decimal places (i.e. 0.0001) to permit the calculation of a response ratio. Instead of replacing zeros with the smallest whole number (1), they replaced them with 0.0001 and subtracted the same from values of 100%. Because lnRR is a ratio of the treatment mean/control, the use of an extremely small denominator results in an immensely inflated response ratio. The same applies if the numerator is extremely small, it produces a hugely inflated negative lnRR. For example, if the control mean is 0% and the treatment is 100%, then: ln(99.9999/0.0001) = 13.8154 using 0.0001 to correct for zero values. By contrast, ln(99/1) = 4.595 using the smallest whole number (1) to correct for zero values. In other words, the estimated response ratio is three times larger when a small fractional value is used to replace zero in percentage data compared with using the smallest whole number in a data range from 0 to 100%. Clements et al.’s decision to replace zeros in percentage data with 0.0001 is especially perplexing when the resolution of the studies involved is considered. Measuring any fish behaviour to an accuracy of 0.0001% would be extraordinarily challenging. Moreover, the original studies in 2009-2010 that reported percentage data [2-4] had a total of 48 observations per trial, therefore, the lowest non-zero value that could be attained (1 in 48) was >2%, which is more than 4 orders of magnitude greater than the 0.0001% non-zero replacement value selected by Clements et al. in their analysis.

The majority of results reported in percentages, and with zero values, are in the first three papers on the topic, published in 2009-2010, so it is no surprise that using 0.0001 to replace zero values throughout the entire data set leads to much larger effect sizes in 2009-2010 compared with subsequent years. This is easily observed in data simulations using either 0.0001, 0.01 or 1 to correct for zero values in percentage data. Using Clements et al.’s data set that has been corrected for data entry errors, screened for inappropriate inclusions (e.g. sham treatments and fluctuating CO2 treatments, see below) and with unexplained missing data sets included (supplementary data https://doi.org/10.25903/d9r4-t979), Fig 2 shows how the decline effect is driven by the choice of replacement values used in percentage and proportional data. When zero values are replaced with 0.0001, the complete, corrected and screened data set exhibits a decline in effect size that is not dissimilar to that originally reported by Clements et al. (2022) (Fig 2a,b), except that the initial decline is less steep (Fig 2c) and the variance-weighted average effect sizes are noticeably higher in 2018-2019 compared with the original (Fig 2d). However, the decline effect (Fig 2e) and the magnitude of weighted average effect sizes (Fig 2f) is markedly reduced if 0.01 is used to correct for zero values in both percentage and proportional data. Most notably, the decline in effect size becomes even flatter (Fig 2g), and weighted effect sizes many times smaller in 2009, 2010 and 2014 (Fig 2h), when zero values in percentage data are replaced with the smallest whole number (1) and replaced with 0.01 for proportional data. From this comparison it is clear to see that Clements et al. claim of an extreme decline effect is a statistical illusion driven by their method of correcting for zero values in percentage data. Indeed, Lajeunesse (2015) [5] warns that “log-ratio effect sizes estimated with RR are at the greatest risk of bias when: (1) the means have small sample sizes, (2) the two means are not close to one another, and (3) at least one of the control and treatment means is near zero” all of which apply to this analysis.

**Figure 2.**
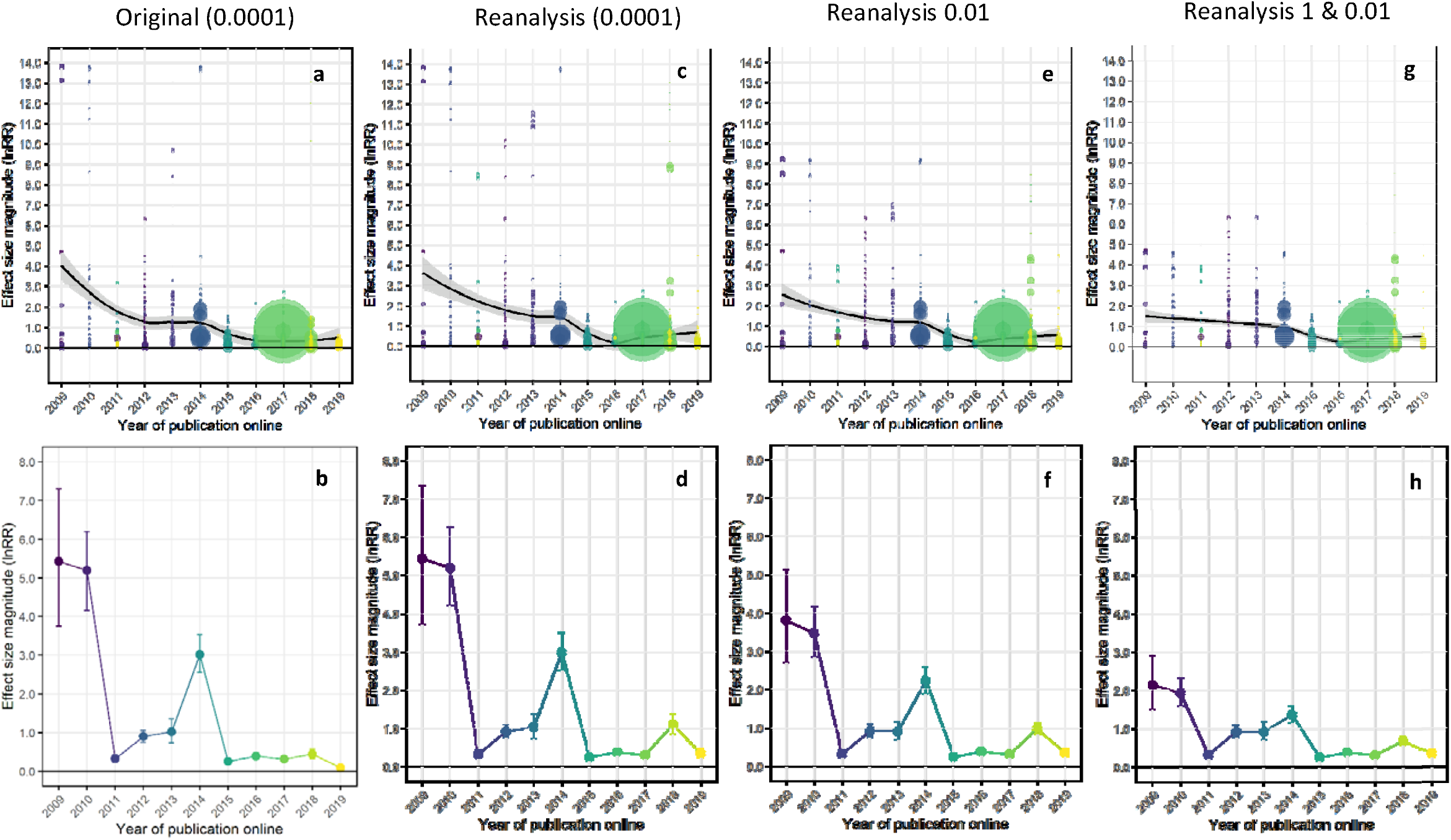
Comparison of decline effect and variance-weighted mean effect sizes with different methods of correcting for zero values in percentage and proportional data. Original data from Clements et al. using 0.0001 (a,b) and reanalysis with corrected, updated and screened data using 0.0001 (c,d), corrected, updated and screened data using 0.01 (e,f) and corrected, updated and screened data using 1 for percentage data and 0.01 for proportional data (g,h).

In addition to the statistical problem associated with correcting for zero values, the analysis by Clements et al. (2022) contains data handling errors, improper data inclusions and exclusions, and inexplicably missing experiments and studies (see table in the methods below), all of which exaggerate the decline effect from the earliest studies and are indicative of biased interpretations.

A preliminary check of the data used in Clements et al. (2022) reveals data entry errors and incorrect values in key treatments (red highlights in supplementary data) that cause effect sizes to be lower than the true value for studies after 2010. For example, the feeding strikes data for McMahon et al. 2018 [7] (study a78) does not match the figure or the underlying raw data in any way whatsoever, and there are errors in the reported N values, despite the correct data being publicly available online since the paper was published. The incorrect data produces much smaller effect sizes for this study than the true values. There are also mistakes in the coding of cue type and life stage of some studies. It is troubling to find such mistakes in a paper that attempts to discredit the research findings of others. Clark, Jutfelt and Sundin have stridently claimed that (unintentional) human errors in data compilation identified in some previous data sets associated with papers from my research group are evidence of data fabrication. Yet, here there are clear data handling errors and incorrect values in the data set used in Clements et al. (2022), as well as incorrect values in the year of publication online and print columns for numerous files (see methods), which was curated and validated by Clements, Sundin, Jutfelt and Clark (see author contributions). Do we conclude this is evidence of research misconduct and data fabrication by these authors? If they apply equal standards to their own work then they must conclude that it is, especially when the errors serve to support the narrative of their paper. These mistakes illustrate how easy it is to make unintentional data handling errors in large, complex data sets, even by authors who have been highly critical of others for doing just that.

A more systemic problem throughout the dataset that leads to artificially diminished effects sizes in papers after 2010 is the inclusion of procedural controls and sham treatments in the calculation of OA treatment effect sizes (blue highlights in supplementary data). By definition, procedural controls and sham treatments are predicted to have no or very small effects if an experiment is working properly. They are designed to check that the experimental method is sound, not to directly test for treatment effects, which in this case is usually the effects of the OA treatment on the behavioural response to a stimulus, such as the presence of a predator or conspecific alarm cues. By including these methodological controls as experiments in their analyses, Clements et al. (2022) have artificially diluted the effect size for several studies conducted after 2010. Furthermore, the 2009-2010 studies did not have procedural controls, whereas procedural controls and shams were used in some studies post 2010. Including methodological controls in the analysis leads to a misrepresentation of the average effects size in some studies and makes it impossible to fairly compare the average effect sizes of papers from 2009-2010 with those published after 2010.

At the same time as including procedural controls and sham treatments that have small or no effects, Clements et al. chose to exclude results where there was a different direction of responses between the control and the OA treatment (i.e. the control might be strongly negative in response to a cue whereas the OA group exhibits a weak positive response to the same cue). The problem here is that these are often the stronger results directly attributable to OA effects, precisely because the treatment effects goes in the opposite direction to the control. For example, the three species for which strong OA effects are observed at 850 ppm CO2 are excluded in the data set for Ferrari et al. 2011 [8] (study a6), leaving only the one species that was found to be much more tolerant of elevated CO2 in the analysis. Similarly, the main results for change in area used were excluded from Ferrari et al. 2012 [9] (study a11). By excluding some of the strongest effects, while retaining weaker effect from the same experiments, Clements et al. (2022) have exacerbated the decline in effect size of experiments immediately after 2010. Moreover, there is a simple solution to the analytical problem of calculating lnRR when there is a different direction of response between the control and OA treatment. One simply needs to replace the small positive number in the OA treatment (or control) with a small negative number (as was done for zero values throughout) to make the calculation possible and retain the strongest OA effects in these studies. This has been done in the data set used here for reanalysis (yellow highlights in supplementary data).

A further issue is the inclusion of treatments that are known to diminish the magnitude of OA effects, such as fluctuating CO2 treatments, that were not included in the original studies. For example, Jarrold et al. 2017 [10] (study a64) showed that daily CO2 cycles greatly diminish the behavioural effects of OA compared with stable elevated CO2 treatments used in earlier studies. By including these treatments in their analysis, Clements et al. diminish the average effects size that would otherwise be attained. Comparing the effects size of experiments with fluctuating CO2 included in the OA treatments, when this was not done in the original studies, is like comparing the effect size of a poison when the antidote has already been administered. It is illogical to include fluctuating CO2 treatments if the aim is to make a fair comparison of effect size through time.

Finally, some experiments and whole studies with strong effects are inexplicably missing in Clements et al.’s data set (e.g. survival data in Davis et al. (2018) [11], study a72), including recent studies by Lecchini et al. (2016) [12], Paula et al. (2019) [13] and Williams et al. (2019) [14] that would be well known to the authors of Clements et al. (2022) (purple highlights in supplementary data). The absence of these studies causes the mean effect size estimated by Clements et al. for studies published in 2018-2019 to be lower than it should be (mean magnitude original vs reanalysis (0.0001) 2018: 0.443 vs 1.111, 2019: 0.088 vs 0.356). Moreover, the mean effect size of studies in 2019 does not fall to zero as reported in Clements et al. (2022) when these studies are included (Fig 1c,d).

Without doubt there has been a decline through time in the averaged effect size from experiments investigating the behavioural effects of OA on fish. This can be seen in the reanalysis of Clements et al. results presented here (Fig 1c), but it is not the extreme decline erroneously reported by Clements et al. (2022). A decline in effect size is not surprising as more and different species are tested, some of which will be much less sensitive to the effects of OA than the orange clownfish, which was the first species tested in this field of study. Indeed, subsequent studies from my own lab show that some other species are unaffected by OA conditions [e.g. 15]. Furthermore, an increasing range of different behaviours have been tested through time, many of which are less affected by OA and generate smaller effect sizes than the initial effects of OA on the response of larvae to concentrated predator odor and habitat cues [e.g. 16]. Methods have also changed through time, in ways that reduce effect sizes compared with the earliest studies in the field [17]. It is not at all surprising there is a decline in the average effect size as more species and different behaviours are tested, and as experiments become more nuanced or include other factors that eradicate or dampen the behavioural impacts of OA [17].

To properly test for a decline effect, it is necessary to compare studies that investigate the same underlying process or mechanism. The pioneering studies into the effects of OA on fish behaviour from 2009-2010 [2-4] tested the olfactory-mediated behavioural response of larval clownfish to predator and habitat cues. By screening the updated data set to consider only studies that examine olfactory-mediated responses to risk cues (predator or conspecific alarm cue) or habitat cues (physical habitat or resident fishes) (supplementary data) it is possible to directly compare the results of the earliest studies with those done after 2010. Contrary to the claims of Clements et al. [1] there is considerable consistency in the effects sizes of the earliest and subsequent studies when just olfactory cues for risk and habitat are considered in the corrected, complete, and properly screened data set (Fig 3a). The average weighted effect sizes for 2009-2010 (mean magnitude 2.149 and 1.939, respectively) overlap with 2012, 2013, 2014 and 2016 (1.461, 1.930, 1.955, 1.1436, 0.939) (Fig 3b) and in 2018 there are experiments with equal effect sizes (lnRR) to those observed in 2009-2010 (Fig 3a). The contrast with Clements et al.’s conclusions could not be starker.

**Figure 3.**
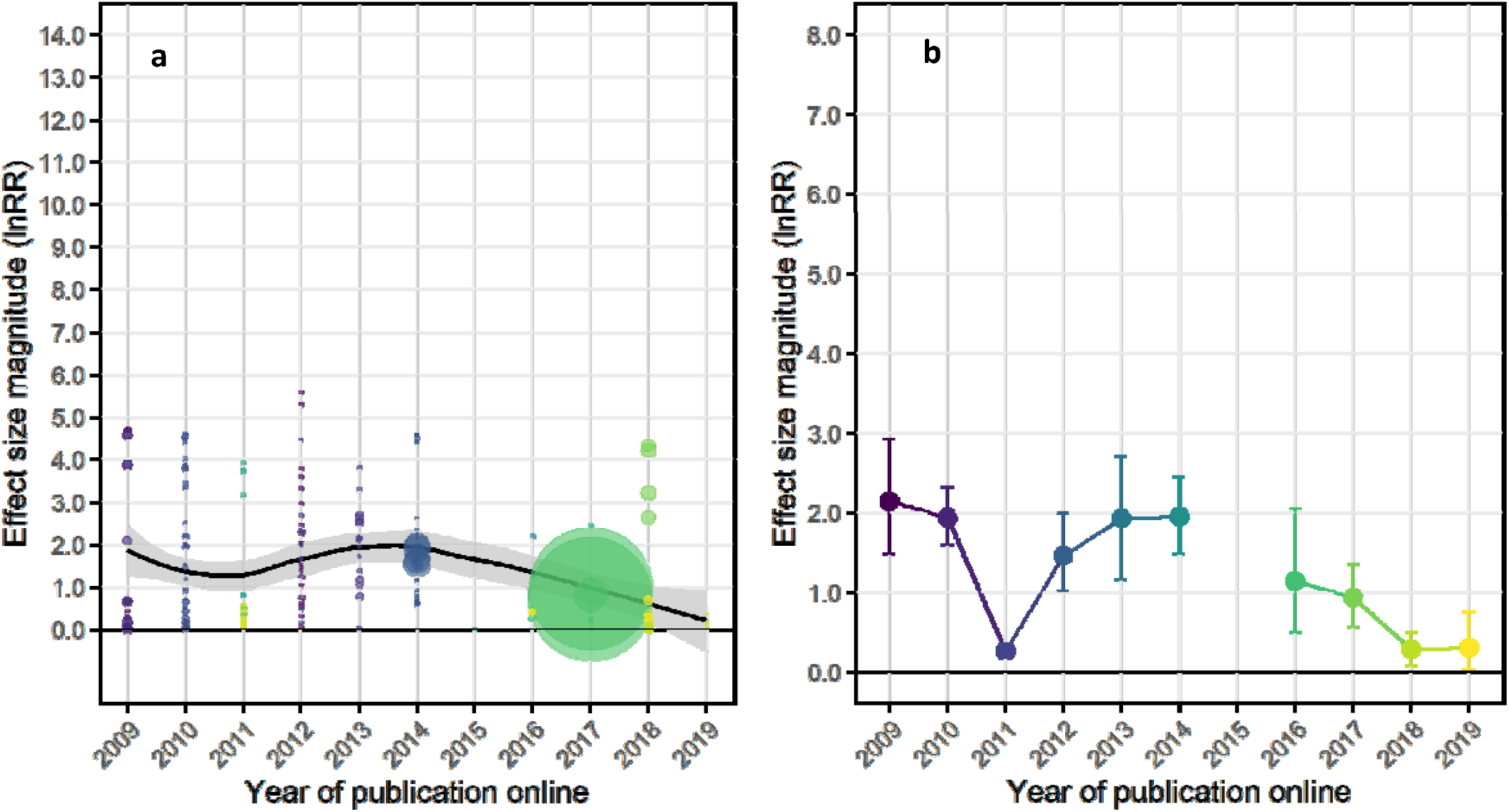
Reanalysis of the decline effect using only studies that examine olfactory-mediated responses to risk cues (predator or conspecific alarm cue) or habitat cues (physical habitat or resident fishes) with the corrected, updated and properly screened data set using 1 for percentage data and 0.01 for proportional data (supplementary data 2). (a) shows unweighted effect sizes (lnRR) fitted with a Loess curve and 95% confidence bounds and (b) shows the modelled variance-weighted average effect sizes by year. There was only one relevant data point in 2015 (study a58), which meant that modelling for Fig 3b was not possible for that year.

Effect size meta-analysis is a useful tool, but it needs to be properly applied and interpreted. Even finding a weak effect averaged across many studies does not necessarily mean that there are not meaningful and important effects of the variable in question. Especially in ecology, environmental impacts on one or a few species or traits can have important consequences on populations, communities and ecosystems. Indeed, recent research, some of which was not captured in Clements et al. analysis [12-14, 18], continues to show that future OA conditions can affect critically important behaviours in coral reef and other fishes. The authors of Clements et al. have been vocal about maintaining the highest standards of research methodology and integrity, yet the extensive problems identified in this paper show they fall well short on the ideals they demand of others.

## Methods

I manually screened Clements et al.’s S2 raw data file for errors, inappropriate inclusions and missing data (listed in the table below). These are highlighted in red, blue and purple in the supplementary data available at Research Data JCU (https://doi.org/10.25903/d9r4-t979). Specific experiments excluded in Clement’s et al.’s S2 raw data file because there was a different direction of responses between the control and the OA treatment were adjusted to enable inclusion by replacing the small number in the OA treatment (or control) with a very small number of the same sign as the control (or OA treatment), as was done by Clements et al. for zero values throughout the data set. These corrections are highlighted in yellow in the supplementary data and listed in the table below.

| Correction type | Highlight | Study | Rows in corrected data file |
| --- | --- | --- | --- |
| Data errors | Red | a4 | 69, 70, 73, 74, 77, 78 |
|  | Red | a23 | 243, 244, 245, 246 |
|  | Red | a78 | 669, 670, 671, 672 |
| Incorrect inclusions | Blue | a11 | 120, 122, 124, 126, 128, 130, 132, 134, 136, 138, 140, 142, 144, 146, 148, 150, 153, 156 |
|  | Blue | a22 | 215, 218, 221, 224, 227, 230 |
|  | Blue | a27 | 285, 286, 289, 290 |
|  | Blue | a64 | 554, 545, 547, 548, 550, 552, 554, 556 |
|  | Blue | a65 | 558, 560, 562, 564, 566, 568 |
|  | Blue | a70 | 601, 602 |
|  | Blue | a78 | 665, 666, 667, 668 |
| Missing data | Purple | a23 | 247, 248, 249, 250, 251, 252 |
|  | Purple | a72 | 616, 617 |
|  | Purple | a92 | 828, 829 |
|  | Purple | a93 | 830, 831, 832, 833, 834, 835 |
|  | Purple | a94 | 836, 837, 838 |
|  | Purple | a95 | 839, 840 |
| Incorrect exclusions | Yellow | a6 | 90, 92, 94 |
|  | Yellow | a11 | 129, 131, 143 |
|  | Yellow | a22 | 229 |

To perform the reanalysis of the data, the relevant sections of the S1 code file (https://journals.plos.org/plosbiology/article?id=10.1371/journal.pbio.3001511#sec021) was copied into a new R markdown file. The relevant original data files were also downloaded (S2, S5, S7 and S10). The S5, S6 and S7 files all have errors in the columns of publication.online and publication.print, such that the model will not run due to a lack of publications within the year 2009 for publication.online. Unfortunately, it was not a matter of the two columns headings being switched and thus the S2 raw data excel file has been used here to recreate appropriate csv files for analysis (recreation of S5). The S1 file contained code for graphing components of Fig 1B but this did not produce a graph that was visually similar, thus changes and additions to aesthetic related code was made. In addition, the S1 file did not contain the code for Fig 1A, so the code for Fig 1B was modified to allow plotting of visually identical figures, but it may contain slight differences. The same R file was used to run all the reanalyses. Notes in the R markdown file identify where additions or changes were made, all available at Research Data JCU (https://doi.org/10.25903/d9r4-t979). For example, in the case of the olfaction reanalysis the year 2015 did not contain enough data so this part of the code was not included.

